# Stick-slip unfolding favors self-association of expanded *HTT* mRNA

**DOI:** 10.1101/2024.05.31.596809

**Authors:** Brett M. O’Brien, Roumita Moulick, Gabriel Jiménez-Avalos, Nandakumar Rajasekaran, Christian M. Kaiser, Sarah A. Woodson

## Abstract

In Huntington’s Disease (HD) and related disorders, expansion of CAG trinucleotide repeats produces a toxic gain of function in affected neurons. Expanded *huntingtin* (exp*HTT*) mRNA forms aggregates that sequester essential RNA binding proteins, dysregulating mRNA processing and translation. The physical basis of RNA aggregation has been difficult to disentangle owing to the heterogeneous structure of the CAG repeats. Here, we probe the folding and unfolding pathways of exp*HTT* mRNA using single-molecule force spectroscopy. Whereas normal *HTT* mRNAs unfold reversibly and cooperatively, exp*HTT* mRNAs with 20 or 40 CAG repeats slip and unravel non-cooperatively at low tension. Slippage of CAG base pairs is punctuated by concerted rearrangement of adjacent CCG trinucleotides, trapping partially folded structures that readily base pair with another RNA strand. We suggest that the conformational entropy of the CAG repeats, combined with stable CCG base pairs, creates a stick-slip behavior that explains the aggregation propensity of exp*HTT* mRNA.

## INTRODUCTION

Huntington’s Disease (HD), a fatal neurodegeneration, is caused by expansion of the CAG repeats located in the first exon of the *huntingtin* (*HTT*) gene, in which disease severity and age of onset strongly correlate with the number of CAG repeats ^1^. Aggregation of the polyglutamine-containing *HTT* protein has been extensively studied ^2^. Emerging evidence, however, suggests that toxicity of the expanded (exp) *HTT* mRNA also contributes to disease ^3, 4,5^. Firstly, RNAs containing expanded CAG tracts form nuclear or perinuclear aggregates ^6, 7, 8^, and transcription is neurotoxic even when the RNA cannot be translated ^9^. Secondly, a genome wide association study (GWAS) revealed that the length of the uninterrupted CAG repeat tract, rather than polyQ tract length, is the primary determinant of HD age of onset ^10, 11^. Furthermore, cellular models of HD have shown that exp*HTT* mRNA self-associates into foci that sequester the splicing regulatory protein, MBNL1, and other RNA-binding proteins ^8, 12, 13, 14, 15^.

Physical properties that promote RNA aggregation and protein sequestration have been difficult to study because the repetitive sequence adopt many structures. RNAs with 12-15 CAG trinucleotide repeats form hairpins containing A-A mismatches ^16, 17^ that may ‘slip’ into alternative alignments of the opposing strands, exposing a variable number of single-stranded triplets. In simulations, these unpaired CAG triplets seed inter-strand base pairs ^18^, explaining why phase separation propensity increases with the number of CAG triplets ^8, 19^.

Sequences flanking the CAG repeats in *HTT* also affect mRNA structure and may contribute to repeat expansion toxicity. Chemical probing experiments showed that 7-9 flanking CCG proline codons in *HTT* mRNA base pair with CAG triplets, forming a composite hairpin at the base of the stem ^13, 20^ (Fig. 1A). The CCG repeats increase the melting temperature of the RNA by ∼15°C compared to pure CAG repeats of identical length ^13^. It is unknown how CAG-CCG base pairs alter interactions between RNA strands.

**Figure 1:**
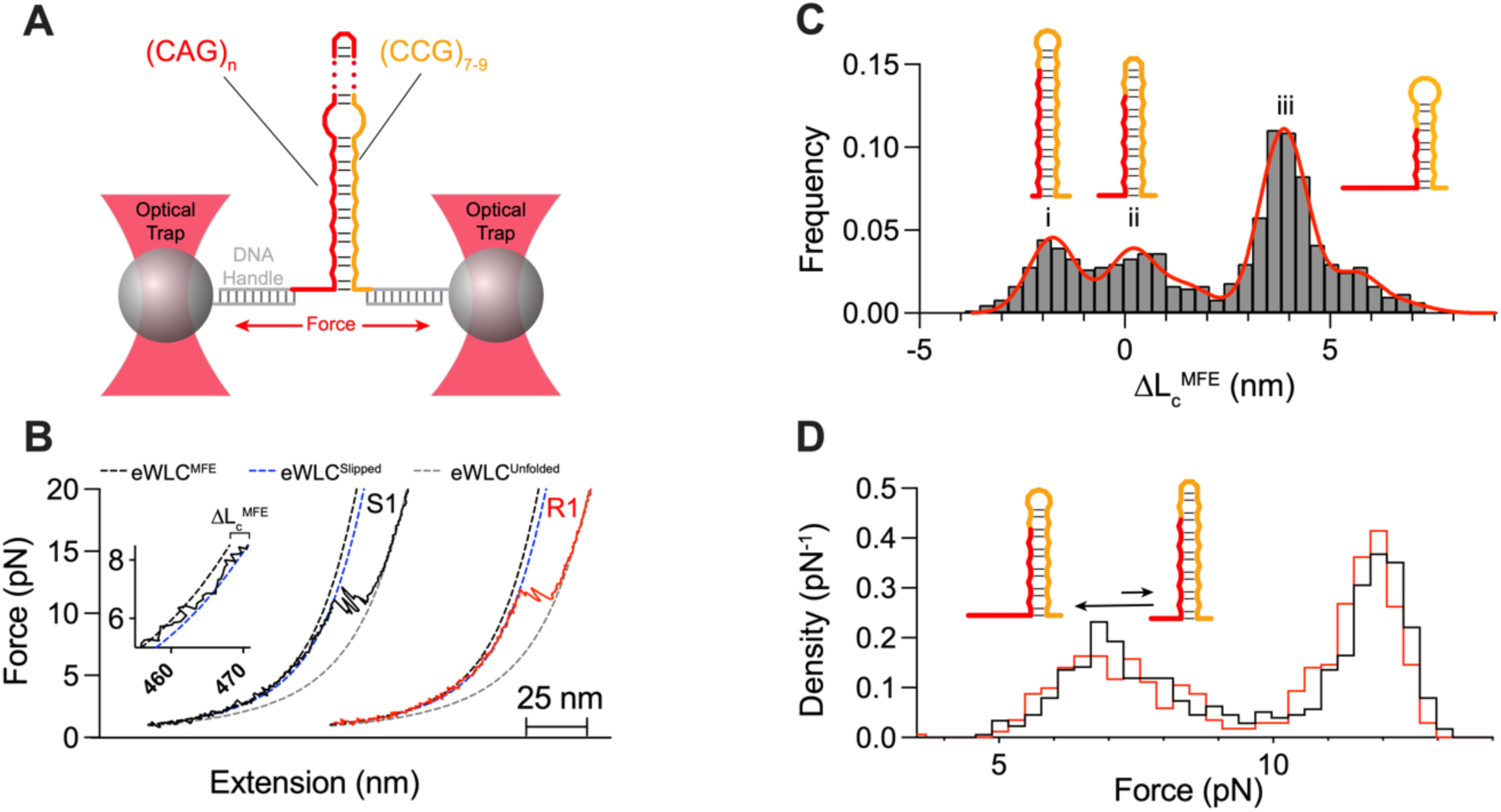
Normal *HTT* RNA with 7 CAG repeats unfolds cooperatively. **A.** RNAs corresponding to the CAG repeat region of human *HTT* exon 1 were attached to optically trapped beads via dsDNA handles. f*HTT*_n_ mRNAs contain *n* = 7, 12, 20 or 40 CAG triplets (red line) plus flanking CCGCAA(CCG)_7_ repeats (gold line). **B.** Example stretch (black, S1) and relax (red, R1) cycles for f*HTT_7_* at 90 nm/s show unfolding/refolding transitions near 11.5 pN. Inset: slip transitions at low tension. Dashed curves represent eWLC models for folded (black), unfolded (gray) and slipped (blue) states. See Fig. S4 for details. S1 and R1 are plotted with an offset for clarity. **C.** Contour lengths of slipped intermediates before the main unfolding transition from eWLC fits (blue dashed line in B) relative to the folded state (eWLC^MFE^ - black dashed line in B). Red line, sum of 6 Gaussians. Peak averages correlate with predicted structures: (i) reference MFE structure with one single-stranded CAG triplet (0.19 ± 0.06 nm), (ii) no unpaired CAG triplet (–1.76 ± 0.04 nm), (iii) three unpaired CAG triplets (3.87 ± 0.02 nm). **D.** Distributions of unfolding (black) and refolding (red) forces, from experiments as in B. (*N* = 286 FECs).

Why neurodegeneration rises around a threshold of 36 CAG repeats, and why some CAG expansion mutations are toxic whereas others are not, remain long-standing questions ^21^. To address these challenges, we probed the folding and unfolding of normal and expanded *HTT* mRNA using single-molecule force spectroscopy with optical tweezers that detect unfolding of a few base pairs at a time ^22, 23^ and resolve heterogeneous folding pathways ^24, 25^. Our results show how continuous, non-cooperative slippage of CAG base pairs hampers complete refolding of exp*HTT* mRNA. ‘Sticky’ interactions with the adjacent CCG trinucleotides stabilize partially slipped structures containing unpaired CAG repeats. The exposed single-strands readily initiate inter-strand base pairing, supporting models in which single-stranded CAG repeats drive phase separation and aggregation of CAG repeat RNA ^18, 26, 27^.

## RESULTS AND DISCUSSION

### Force spectroscopy resolves *HTT* RNA folding intermediates

To investigate *HTT* mRNA structures by force spectroscopy, fragments of human *HTT* exon 1 (f*HTT*) RNA spanning the CAG repeat region were joined with dsDNA handles and attached to polystyrene beads held by optical traps (Fig. 1A). The mRNA fragments contained 7- 40 CAG codons, the adjacent CCG-CAA-(CCG)_7_ codons, and an additional 20 nt *HTT* sequence flanking the trinucleotide repeats (Supplementary Fig. S1 and Table S1). Tethered f*HTT* mRNAs were repeatedly stretched and relaxed at a constant speed, yielding force extension curves (FECs) that revealed the size and mechanical stability of the mRNA structures (Fig. 1B). Fitting an extensible worm-like chain (eWLC) model to the data yielded the changes in contour length (*L_c_*) upon unfolding and refolding (see Methods and Fig. 1B), the forces at which unfolding and refolding occur (see Methods and Supplementary Fig. S2) and the cooperativity of each transition (Supplementary Fig. S3).

mRNA containing 7 CAG triplets (f*HTT*_7_), within the non-pathogenic range, exhibited near-equilibrium, cooperative unfolding and refolding transitions at 11.8 ± 0.56 pN (Fig. 1B). However, the distribution of contour length changes associated with these transitions ranged from 18.2 nm to 29.7 nm compared to 25.96 nm expected for unfolding of the predicted minimal free energy (MFE) hairpin. This difference is explained by smaller transitions below ∼ 7 pN (Fig. 1B inset), indicating that f*HTT*_7_ mRNA populates ‘slipped’ structures before unfolding completely at higher force.

To evaluate these slipped structures, we fitted each FEC with a series of eWLC models to determine the contour length change relative to *L*_c_ of the predicted MFE structure (Δ*L_c_*^MFE^) which is comprised of base paired CAG-CCG triplets plus one unpaired CAG (Supplementary Fig. S4). The distribution of Δ*L_c_*^MFE^ at low tension peaked at values that differed by multiples of one single-stranded trinucleotide (∼1.77 nm; Fig. 1C and Table S2). This length change agreed well with ‘slipped’ conformations of both shorter and longer contour lengths than the predicted MFE structure (Fig. 1C). Control experiments showed that these low force transitions did not arise from transitions in the *HTT* nucleotides flanking the repeat region (Supplementary Fig. S5). The FECs were not well described by sequential opening of the hairpin, indicating that the CAG/CCG repeat hairpin slips rather than partially unfolds at low forces.

The presence of slipped intermediates before the main unfolding transition was supported by two populations of the unfolding and refolding forces (Fig. 1D): transitions centered around 11.8 ± 0.56 pN were almost exclusively associated with cooperative unfolding of the f*HTT*_7_ hairpin, whereas transitions at ∼ 6.9 ± 1 pN corresponded to ‘slippage’ of one to four CAG repeats.

Thus, an unexpanded *HTT* mRNA folds cooperatively into a hairpin that is stabilized by base pairs between CAG and CCG triplets. Yet, even this short mRNA samples different structures resulting from slippage of base pairs within the repeat region. These results agree well with previous single-molecule FRET studies of slippage in DNA hairpins containing pure CAG or CTG repeats ^28, 29, 30^, suggesting that such interconversion is an intrinsic property of nucleic acids containing triplet repeats.

### Non-cooperative unfolding of expanded CAG repeats

Expanding the repeat region beyond seven CAG triplets resulted in progressive loss of large cooperative transitions during hairpin opening and closing (Fig. 2). f*HTT*_12_, containing 12 CAG triplets, visited a greater number of slipped intermediates than f*HTT*_7_, but still unfolded cooperatively at 12.3 ± 0.68 pN (Fig. 2A, first row). By contrast, expansion to 20 or 40 CAG triplets caused a marked reduction of folding cooperativity (Fig. 2A, second and third rows). Almost half of the FECs for f*HTT*_20_ and nearly all FECs for f*HTT*_40_ showed gradual non-cooperative extension between 8.5-11 pN. Analogous non-cooperative unfolding was observed for a tetratricopeptide repeat protein ^31^. As the mRNA structure unraveled, transitions to structures that were slightly more or less extended than the preceding conformation indicated reorganization of the base pairs.

**Figure 2:**
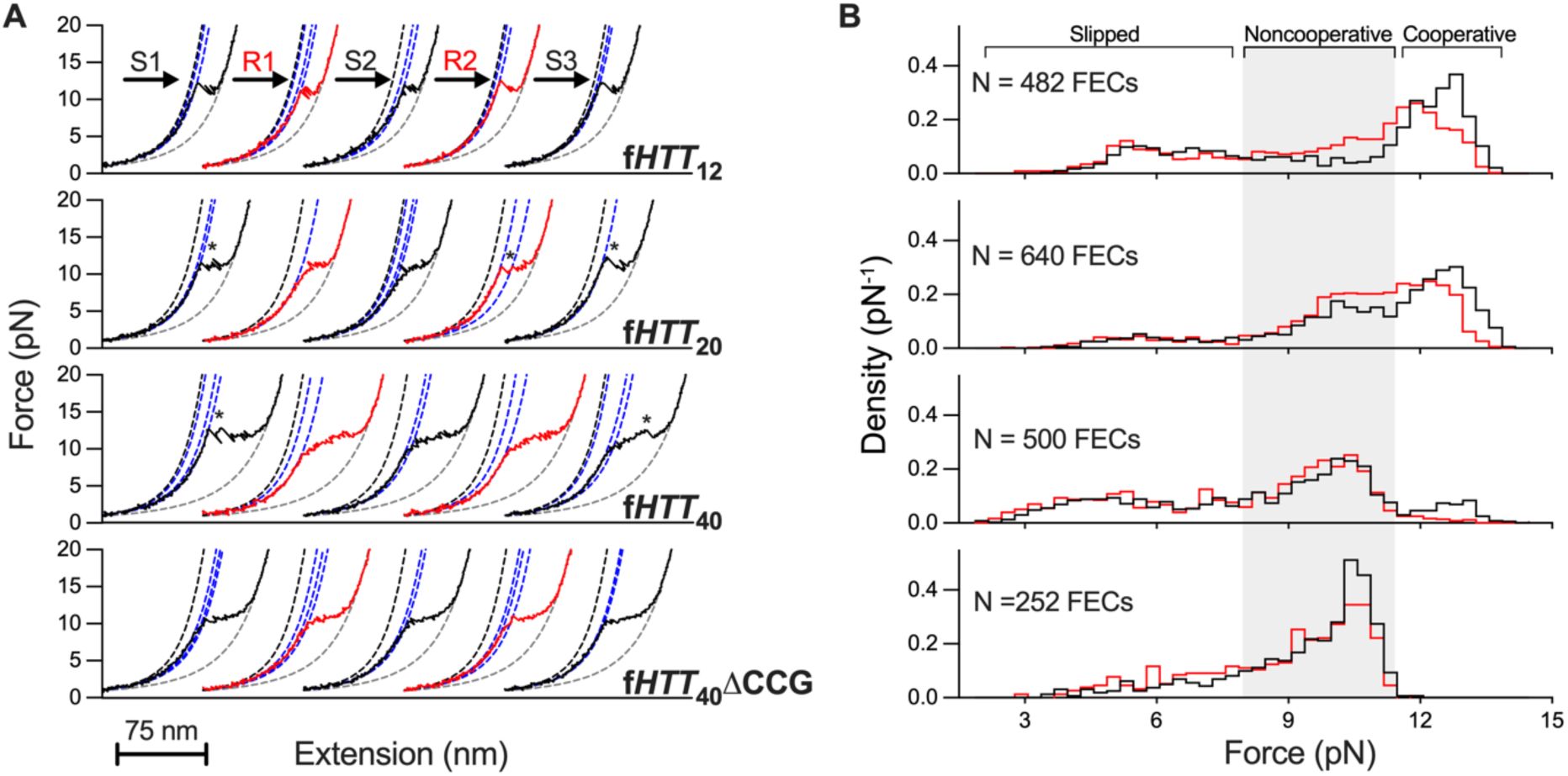
Expanded HTT mRNA unravels non-cooperatively. **A.** Three successive stretch (black) and relaxation (red) cycles for f*HTT*_12-40_ mRNAs, fit to eWLC models as in Fig. 1. Cooperative transitions (asterisks) become less common as the repeat expands and are lost for ΔCCG (bottom). **B.** Probability density of unfolding (black) and refolding (red) forces for each f*HTT* mRNA. Average unfolding force for fHTT_12_ was 12.4 pN, compared to 11.8 pN for fHTT_7_. Shaded region indicates non-cooperative transitions.

Due to hairpin slippage, non-cooperative unfolding of the repeat region occurred at lower forces than cooperative unfolding (8.5-11 pN vs. 12-12.5 pN, respectively; Fig. 2B), and the unfolding force remained constant as the CAG repeats expanded from 12 to 40. Initial and subsequent stretches of f*HTT*_40_ populated similar slipped intermediates (Supplementary Fig. 6A). However, cooperative unfolding of f*HTT*_40_ at high force was observed more frequently during the first stretch of a molecule (Fig. 2A and Supplementary Fig. 6B,C), indicating that the long CAG hairpin did not fully equilibrate at low tension. The fraction of cooperative transitions remained unchanged when the equilibration between force-ramp cycles was increased to 10 s (Supplementary Fig. 6D), a time that is sufficient to equilibrate many RNA hairpins at room temperature ^32, 33, 34^.

Altogether, these results suggested that non-cooperative unraveling and contraction of the CAG region arises from frequent slippage of base pairs that becomes more prevalent as the repeats expand. As the hairpins slip, exposed single-stranded bases pair again in a new register, reorganizing the structure (Supplementary Fig. S7). Applied force favors slippage and thus reorganization, explaining why we observe jumps to both shorter and longer structures as the tension rises. When relaxing the force, random nucleation of CAG hairpins leads to kinetically trapped, misaligned hairpins that unfold non-cooperatively during the subsequent stretch.

### Flanking CCG repeats introduce ‘sticky’ interactions

We next asked what other features of f*HTT* mRNA contribute to its rugged folding landscape. A comparison of successive stretch and relaxation cycles of f*HTT*_40_ mRNA showed that the unfolding path varied between pulls on the same molecule (Fig. 2A). Gradual extension of the repeat region was often punctuated by discrete unfolding transitions or rapid hopping between structures before the mRNA was fully unfolded (Fig. 2A, third row, last FEC). This variation contributed to the broad distribution of intermediates and unfolding forces.

As cooperative transitions occurred at higher forces than non-cooperative transitions (Fig. 2B), they must reflect the unfolding of mechanically stable structures. We conjectured that these stable structures arise from interactions with the flanking CCG repeats, which are known to increase the mRNA melting temperature ^13^. Owing to sequence degeneracy, base pairing of the CCG repeats with any of the 40 CAG repeats can give rise to many intermediate structures.

To test this possibility, we measured the folding of f*HTT*_40_ΔCCG mRNA containing only 40 CAG repeats but no CCG repeats. Deletion of the flanking CCG repeats eliminated the cooperative transitions, resulting in smooth non-cooperative unraveling (Fig. 2A, bottom). The elimination of cooperative transitions correlated with a narrower distribution of unfolding forces and fewer resolved slipped intermediates for f*HTT*_40_ΔCCG than for f*HTT*_12_, f*HTT*_20_ or f*HTT*_40_ (Fig. 2B, bottom).

We concluded that the cooperative transitions observed in f*HTT* mRNAs correspond to folding and unfolding of hairpins with CAG-CCG base pairs at the base of the stem to which tension is being applied. Base pairing between CAG and CCG triplets at the ends of the repeat region becomes less likely as the number of CAG triplets increases, explaining the observed decline in large cooperative transitions (Fig. 2B). However, locally stable structures created by CCG repeats can unfold at any point during unraveling of the repeat region. Therefore, we propose that the more stable base pairs formed by the flanking CCG repeats create ‘sticky’ interactions that hinder the facile slippage of CAG-CAG base pairs within expanded *HTT* mRNA.

### Variable refolding in expanded *HTT* mRNAs

Inspection of individual FECs showed that f*HTT*_40_ RNA traveled through many intermediate conformations at low force before unfolding at high force (Fig. 3A). To quantify the occupancy of slipped conformations, we calculated the contour lengths of all identifiable intermediates before the high force unfolding transition as in Fig. 1C (Methods). As anticipated, the distribution of intermediates with different Δ*L_c_*^MFE^ values broadened as the number of CAG triplets expanded from 7 to 40 (Fig. 3B). Whereas f*HTT*_7_ populated a few intermediates with up to four unpaired CAG triplets, f*HTT*_12_ occupied a wider distribution of intermediates that shifted to even more extended conformations for f*HTT*_20_ and f*HTT*_40_ (Fig. 3B). Distinct peaks in the Δ*L_c_*^MFE^ distribution separated by Δ*L*_c_ ∼ 2 CAG repeats suggested a preference for folding steps involving an even-number of CAG triplets, similar to CAG DNA hairpins ^28^.

**Figure 3.**
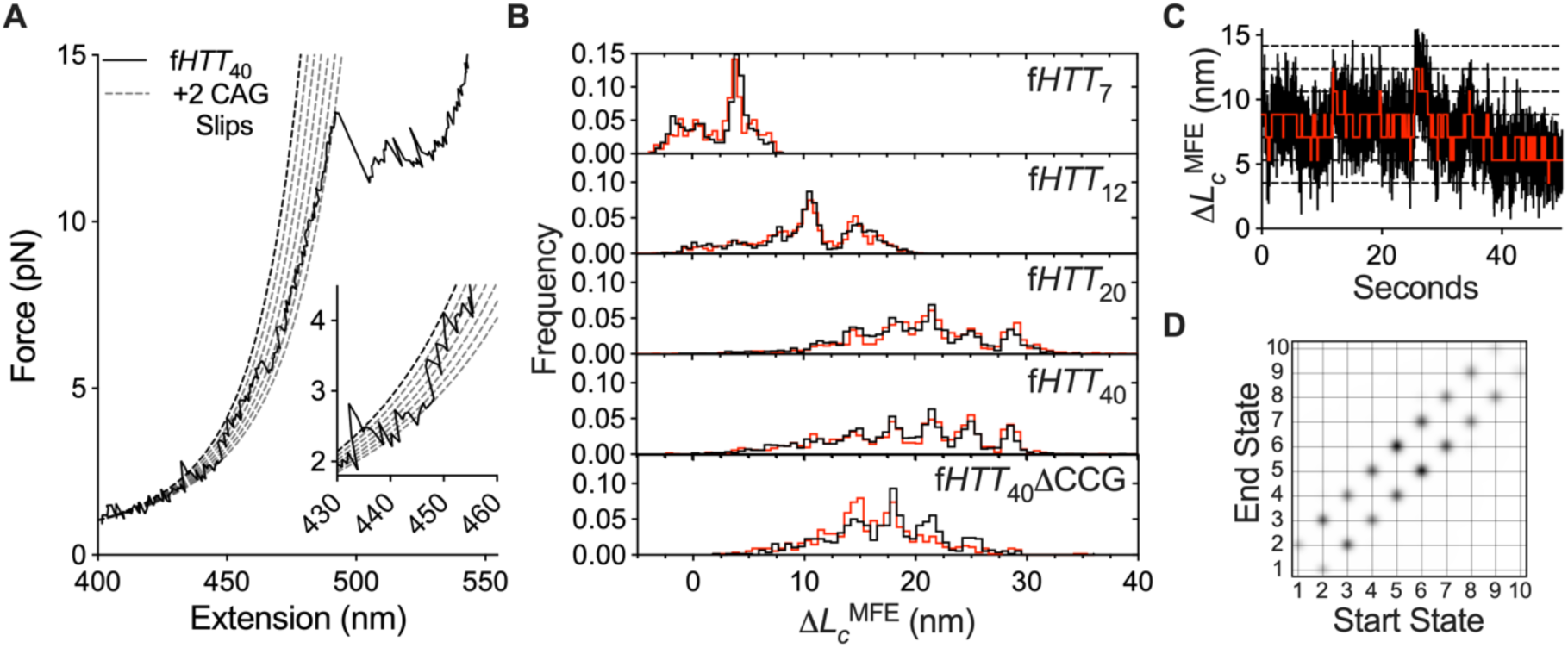
Continuous stepwise slippage of expanded CAG tract. Slipped intermediates of f*HTT*_40_ analyzed as in Fig. 1. **A.** Example FEC demonstrating transitions at low force that extend or shorten *L*_c_, before unfolding at 13.2 pN; see Supplementary Figs. 2-3 for analysis details. Dashed lines, eWLC fits to structures differing by 2 unpaired CAG triplets. Inset: Expansion of low force region. **B**. Distribution of Δ*L_c_*^MFE^ for all f*HTT* mRNAs. Stretch, black; relax; red. **C.** Passive mode fluctuations in Δ*L_c_*^MFE^ for f*HTT*_40_ at ∼ 6 pN pretension (red circle in **A**). Black dashed lines show states used to fit HMM model, red line shows HMM fit. Similar results were obtained when the HMM fit was unconstrained (Fig. S10). Hopping rates of 0.9-3 s^-1^ are comparable to the folding kinetics of a stable 21 bp hairpin ^32, 33^. **D.** Transition density plot from the HMM model in C. See Fig. S11 for data at high force.

Deletion of the flanking CCG repeats narrowed the distribution of intermediates and shifted the population to shorter Δ*L_c_*^MFE^ values, suggesting that pure CAG repeats refold into a narrower range of structures than f*HTT*_40_ (Fig. 3B, bottom). This observation supported our conclusion that the CCG repeats impede relaxation of the mRNA structures and increase the number of structures formed.

### Conformational entropy of exp*HTT* mRNA is inherent to its sequence

Our results indicated that the degeneracy of base pairing configurations hinder exp*HTT* mRNA from adopting any single structure. This interpretation was supported by the similar range of contour lengths sampled during the first and subsequent stretches of f*HTT*_40_ mRNA (Supplementary Fig. S6A). These intermediates were not due to differences between molecules, as different f*HTT*_40_ molecules sample similar distributions of *L*_c_ and *F*_U_ (Supplementary Fig. S8). Similar folding intermediates were observed in NaCl or KCl (Supplementary Fig. S9). Extended intermediates were more prevalent in Mg^2+^ or upon rapid release of tension (‘snap relax’; Supplementary Fig. S9), presumably because partially folded structures are more easily trapped. This structural heterogeneity observed over a range of conditions suggested that high conformational entropy is an intrinsic feature of exp*HTT* mRNA.

### Slippage occurs in sequential steps

To investigate how f*HTT*_40_ mRNA dynamically slips between different secondary structures, we performed passive mode experiments in which the optical traps were kept at a constant separation ^32^, revealing slippage events with discrete changes in force over time (Fig. 3C, Methods). Changes in force were converted into Δ*L_c_*^MFE^ and a hidden Markov model (HMM) was used to obtain the transition probabilities (Fig. 3C, Methods). At low force (∼ 5-9 pN), the data were best described by ten states, with force-dependent dwell times ranging from 0.3 – 1.1 s (Table S3). Each state differed by the contour length of one CAG trinucleotide (Supplementary Fig. S10). The transition density plot (TDP, Fig. 3D) revealed that most transitions occur between adjacent slipped conformations. These measurements established slippage by one trinucleotide as the fundamental unit of exp*HTT* conformational dynamics.

When f*HTT*_40_ was held at a higher pre-tension near the global unfolding transition, hopping between conformations of similar Δ*L_c_*^MFE^ was punctuated by brief excursions to more folded states (Supplementary Fig. S11, Table S4). These unexpected transitions to shorter structures were associated with varied changes in Δ*L_c_*^MFE^. This variation can be explained by transient base pairing between the CCG repeats and different segments of the CAG tract that become exposed at high force. Thus, the force-ramp and passive mode results show that the expanded array of CAG repeats hops or slips between different base pairing registers that expose or sequester CAG trinucleotides.

### Slipped CAG base pairs enable inter-strand pairing

We lastly asked whether the dynamic rearrangements observed for f*HTT*_40_ could facilitate exp*HTT* mRNA aggregation. Unpaired CAG triplets exposed by non-cooperative slippage could serve as toeholds for intermolecular base pairing, which is the first step of RNA aggregation ^13,18^. To test this idea, we moved a single tethered f*HTT*_40_ mRNA into a microfluidic channel containing 500 nM free f*HTT*_40_ mRNA (Fig. 4A). Strikingly, we immediately observed new features in the FEC, compared to the preceding force-ramp cycles of the same tethered mRNA in buffer only (Fig. 4B, top). Because these features were never observed for isolated mRNA, they must be a direct result of inter-strand interactions between tethered and freely diffusing f*HTT*_40_.

**Figure 4.**
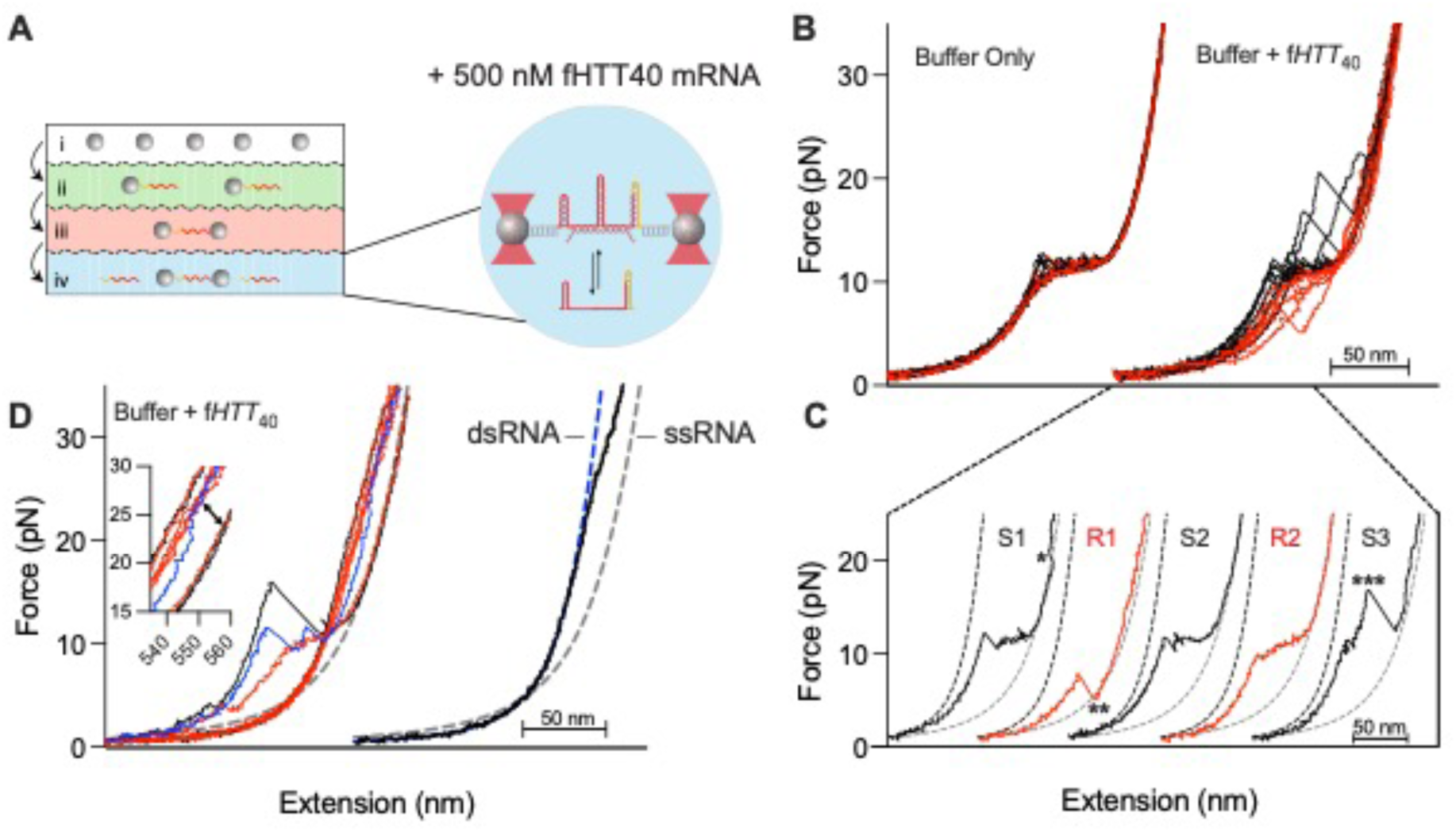
Intermolecular association of partially unfolded exp*HTT* RNA. **A.** Tethered f*HTT*_40_ mRNA is moved from a microfluidic channel containing buffer (i-iii) to one containing 500 nM free f*HTT*_40_ mRNA (iv), where it can form intermolecular base pairs. **B.** Example stretch (black) and relax (red) FECs in buffer (left) and for the same molecule after exposure to free mRNA (right). **C**. Successive cycles showing (*) free RNA binding at high force; (**) release at low force; (***) unfolding and partial release at high force. **D.** Conversion to dsRNA after several cycles. Black, last full extension; red, relaxation; blue, converts to dsRNA at high force (inset) and remains base paired during subsequent cycles. Right, observed FEC compared with dsRNA eWLC model (blue dashed line).

In one example, the first extension in the presence of free mRNA was normal up to ∼18 pN, at which point the unfolded tethered mRNA jumped to a shorter structure (Fig. 4B inset, FEC S1). Upon relaxation of the force, the mRNA did not refold gradually around 10 pN as usual, but instead refolded in a single step at 7.5 pN (Fig. 4B inset, FEC R1). The next force-ramp cycle displayed the expected force-extension behavior (Fig. 4B inset, FECs S2, R2). These perturbations can be explained by partial inter-strand base pairing with a diffusing mRNA at high force that prevents the tethered mRNA from refolding, until dissociation of the diffusing mRNA at 5 pN allows the tethered f*HTT*_40_ to reestablish intra-strand base pairs.

In another example, the tethered f*HTT*_40_ remained folded until abruptly opening at ∼ 17 pN, far above its usual unfolding force (Fig. 4B FEC S3). Unfolding at high force can be explained by inter-strand base pairing that bridges the CAG tract like a cruciform, preventing the tethered RNA from unfolding until one side of the complex dissociates. Interactions between RNA molecules typically occurred near the unfolding force for the tethered RNA, indicating that partial unfolding of the tethered RNA favors inter-strand base pairing. Exposed CAG triplets in the free RNA likely also contribute.

After a few force-ramp cycles in the presence of free RNA, several f*HTT*_40_ molecules failed to refold, and remained unfolded during subsequent stretch-relax cycles (Fig. 4C, left). The FECs for these molecules were described well by the eWLC of an RNA duplex the length of f*HTT*_40_, with a persistence length of 40 nm (Fig. 4C, right). Therefore, inter-strand base pairing, once initiated, can persist and ultimately replace the self-structure of the tethered RNA.

## CONCLUSIONS

Our single-molecule force spectroscopy results reveal an unanticipated dynamical transition in exp*HTT* mRNA at the pathogenic threshold of ≥36 CAG repeats ^1^ ^35^. Short f*HTT* mRNAs with 7 or 12 CAG repeats unfold and refold cooperatively, indicating they form defined hairpins, as previously proposed ^13^ (Fig. 5). By contrast, f*HTT* mRNAs with 20 and especially 40 CAG repeats extend and retract non-cooperatively. This accordion-like unraveling and refolding of the expanded CAG tract arises from constant slippage and reorganization of CAG base pairs, and from heterogeneous nucleation of new hairpins (Fig. 5) ^36^. As a result, the expanded repeats never refold completely. Importantly, the CCG repeats adjacent to the CAG tract in *HTT* increase folding heterogeneity by forming stable base pairs with many segments of the expanded CAG tract, producing kinetically trapped structures. We show that the ‘stick-slip’ behavior arising from a mix of CAG-CAG and CAG-CCG base pairs in exp*HTT* mRNA favors extended structures (Fig. 3) that readily interact with other exp*HTT* mRNA molecules (Fig. 4).

**Figure 5.**
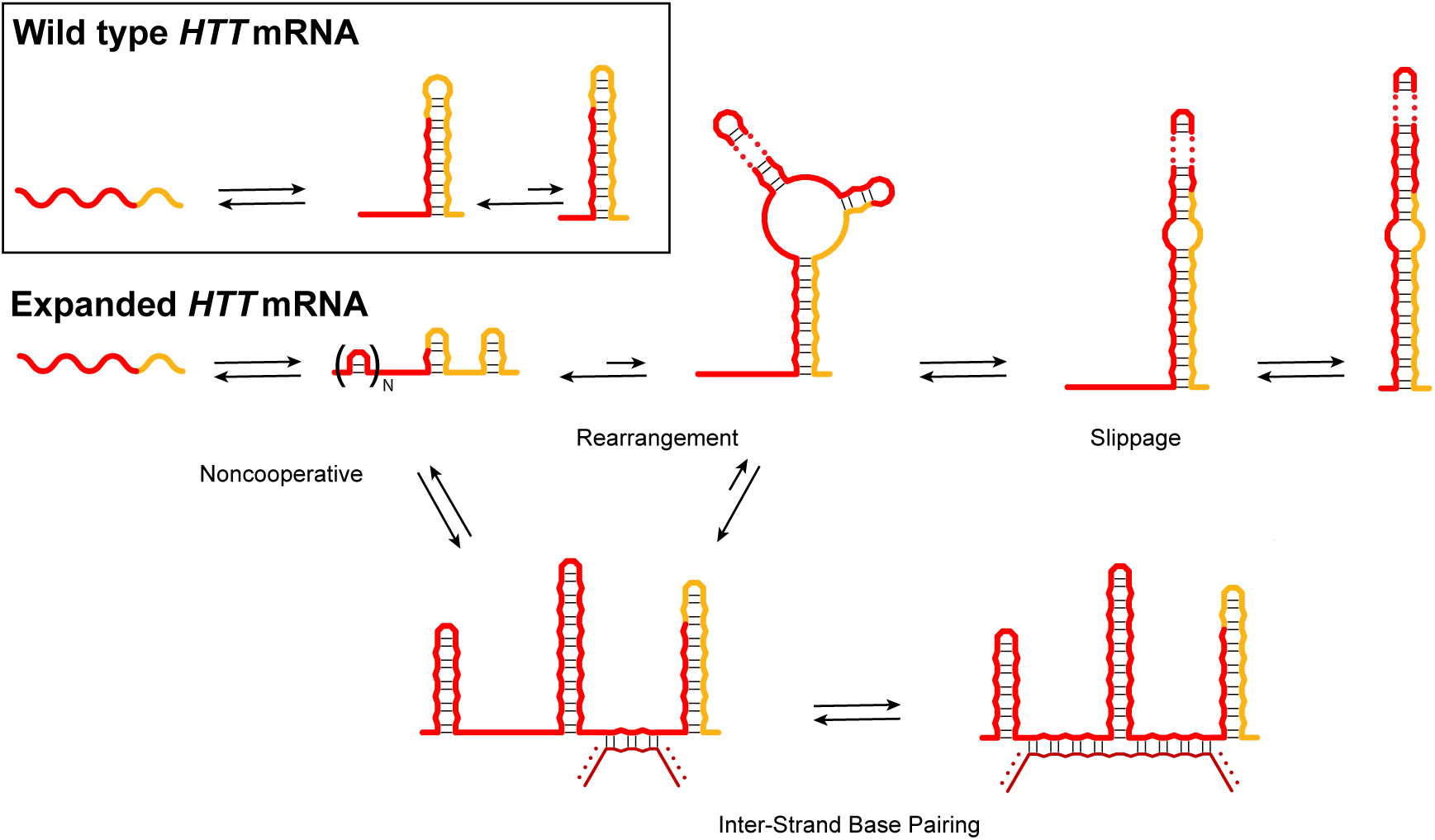
Stick-slip model for CAG repeat aggregation in HD. Short CAG tracts form composite hairpins that unfold and refold cooperatively with limited slippage of base pairs. Expanded CAG repeats unravels and refolds non-cooperatively like an accordion, producing a mixture of secondary structures that include unstable, slippery CAG hairpins and stable, sticky CAG•CCG hairpins that unfold in small rips, producing single-stranded regions that nucleate base pairing with another RNA, producing a network that can lead to aggregation (bottom). Although the intermediate structures vary, the cartoons illustrate conformations that agree with the force extension results.

In our force-ramp (Fig. 2) and passive mode experiments (Fig. 3), reorganization of the CAG tract is stimulated by applied force, consistent with destabilization of one or more base pairs during the transitions between slipped conformations. Slippage could arise from end fraying ^26^ or the propagation of internal bubbles, as recently proposed for DNA repeats ^30^. Alternatively, the two strands of a CAG hairpin may slide past each other via continuous exchange of hydrogen bonds between neighboring nucleobases. This type of exchange does not interrupt base stacking and can occur quickly in double-stranded RNA ^37^ and DNA homopolymers ^38^.

It has been recently suggested that the capacity of repeat RNAs to form heterogeneous interactions drives phase separation into droplets or aggregates ^18, 27, 39^ that have been linked to neurotoxicity ^9, 13, 40^. We observe that the mixed CAG/CCG repeats present in *HTT* mRNA produce a ‘stick-slip’ behavior that we propose makes mixed repeats more prone to condensation and aggregation than pure CAG repeats. We note that HD is one of at least two trinucleotide repeat expansion disorders in which flanking sequences interact with the CAG repeats ^27^. Our results also explain why ribosomes that exert ∼13 pN on the mRNA ^41^ readily pass through expanded CAG tracts, since unraveling exp*HTT* mRNA (10.0 ± 0.7pN) requires less force than cooperatively unfolding shorter hairpins. However, transient unfolding by ribosomes or helicases is unlikely to dissolve repeat RNA condensates because the network of RNA interactions remodels without complete strand dissociation, as observed in our experiments. The single-molecule platform described here will be useful for probing the effects of proteins or small molecules on folding and aggregation of repeat-containing RNA.

## METHODS

### Sample Preparation

Double-stranded (ds) DNA handles with single-stranded overhangs were prepared via PCR amplification of pROEX_HTa ^42^. Primers used to generate the dsDNA handles are listed in Supplementary Table 1. A 691 bp biotinylated DNA handle with a 29 nt 5′ overhang was amplified by Q5 DNA polymerase (NEB, Cat# M0491S) using a 5′ biotinylated forward primer and a reverse primer containing an abasic site ^43^. A 673 bp digoxigenin-labeled DNA handle was amplified in Q5U PCR master mix (NEB, Cat# M0597L) with a forward primer containing a deoxyuridine and a reverse primer with a 5′ digoxigenin tag. A 30 nt 3′ overhang was created by treating the purified PCR product with USER enzyme mix (NEB, Cat# M5505S) for 15 min at 37°C. All PCR products were cleaned up (Qiagen, Cat# 28104) prior to use.

Transcription templates for fragments of the human *huntingtin* exon 1 (f*HTT*N) were synthesized and cloned into pUC-GW-Amp (Genewiz) to yield pUC-GW-f*HTT*N, in which N designates the number of CAG triplets. Transcription templates contained a T7 promoter, 29 bp complementary to the 5′ overhang of the biotinylated DNA handle, 18 bp native exon 1 sequence, N CAG triplets, CCGCAA(CCG)_7_ (flanking polymorphic repeats), a further 20 bp of native exon 1 sequence downstream of the repeats, 30 bp complementary to the digoxigenin-labeled DNA handle and a T7 terminator. Predicted secondary structures of f*HTT* mRNA fragments are shown in Supplementary Figure S1. Transcription templates were prepared by PCR using primers complementary to the T7 promoter and terminator sequences (see Supplementary Table 1) or by digestion of the plasmid DNA with SphI (NEB, Cat# R3182S). mRNA fragments were prepared by in vitro transcription with T7 RNA polymerase using a ratio of 10:30 AU:GC NTPs. The desired transcripts were purified on denaturing 6-8% polyacrylamide gels.

Purified f*HTT* mRNA transcripts were combined with the dsDNA handles at 25 nM equimolar ratio, and annealed and refolded by denaturation at 85°C for 10 min, 65°C for 1.5 h, 55°C for 1.5 h and slow cooling to 4°C (0.1 °C/s) in 200 mM NaCl, 20 mM PIPES, pH 6.5, 60% formamide ^44^. Formation of the desired DNA-RNA complexes were verified by agarose gel electrophoresis (Supplementary Figure S1), purified using a PCR clean up kit (NEB, Cat# T1030L) and eluted in water. Freshly prepared refolded/annealed complexes (1 µL 25 nM stock; ∼ 2.5 nM final) were combined with 4 µL 0.1% w/v 2.1 µm diameter polystyrene beads coated with anti-digoxigenin antibodies (Spherotech, Cat# DIGP-20-2), diluted to 10 µL with assay buffer (10 mM MOPS, pH 7, 250 mM NaCl, 1 mM EDTA), and incubated for 10 min at room temperature. The bead mixture was then diluted ∼50 fold in assay buffer spiked with RNase inhibitor. 1 µL 0.5% w/v streptavidin coated 2.1 µm diameter polystyrene beads (Spherotech, Cat# SVP-20-5) were diluted into 1 mL assay buffer. The two bead mixtures were loaded into separate channels of the Lumicks C-Trap microfluidics system.

### Force-ramp data collection and analysis

Force-ramp assays were performed using a Lumicks C-Trap instrument with BlueLake software (Lumicks). Briefly, streptavidin and anti-digoxigenin coated beads were collected by two optical traps and brought to a separate channel containing assay buffer spiked with RNase inhibitor, 10 mM sodium azide and protocatechuic acid/protocatechuate-3,4-dioxygenase (PCA/PCD; ^45^) for oxygen scavenging. The optical traps were calibrated to obtain the trap stiffness for each pair of beads. Single molecules were tethered between two beads by slowly bringing the trapped beads together until a small increase in force was observed. Once a tether was made, the optical traps were separated at a constant velocity of 90 nm/s to unfold the mRNA and brought together at the same speed to refold the mRNA. At this pulling speed, the loading rate was estimated to be ∼ 10 pN/s just before the start of the main transition of the force vs time traces. After relaxation, the trap movement was paused for 1 s before repeating the stretch and relaxation cycle. For each f*HTT* mRNA, 20 or more molecules were tethered, and 1-100 force-ramp cycles were performed on each tether for each experiment.

### Force-ramp data analysis

For each bead pair, a force baseline was collected and subtracted from valid force extension curves (FECs). Data were collected at 75 kHz and down-sampled to 100 Hz for analysis and the molecular extension for each tether was calculated using the Lumicks Pylake Python package. Processed FECs were obtained from the moving bead, rather than the static bead. In a comparison set, analysis of processed FECs obtained from the static bead were nearly identical (data not shown). Processed FECs were fit by the extensible worm-like chain (eWLC) model

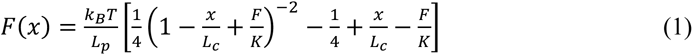

in which *L*_c_ is the contour length, *L*_p_ is the persistence length and *K* is the stretch modulus ^46^. Extension of the folded and unfolded states were modeled by a series of eWLCs describing extension of the dsDNA handles, the ssRNA flanking the CAG hairpin(s) in the folded mRNA, and opening of the mRNA secondary structure. *L*_c_ of the dsDNA handles was fixed at 473.6 nm based on the crystallographic contour length of dsDNA at 0.34 nm/bp, and *L_p_* and *K* were fixed at 40 nm and 1000 pN, respectively, from reported values ^23^. For the ssRNA segments, we used *L_p_* = 1 nm and *K* = 1,500 pN, leaving the *L*_c_^ssRNA^ as the only free fitted parameter ^23^. Extension of the unfolded f*HTT* mRNA was modeled by complete opening of the predicted hairpin structure at 0.59 nm/nt. The unfolded region of each FEC was aligned horizontally (x) to the high force region of the WLC model of the fully unfolded f*HTT* mRNA, followed by a vertical alignment (F) to the low force region of FECs and the WLC for the handles + folded f*HTT* mRNA as previously described ^24^. All analyses were performed on aligned FECs.

The Δ*L_c_* associated with complete opening of the mRNA structure at high force was determined by fitting the region of the FEC just before and after the transition to two eWLC models in series, one for the dsDNA handles and another to describe the extension of the *HTT* ssRNA flanking the folded hairpin. Small changes in contour length below 9 pN were quantified by fitting intermediate regions of FECs with two eWLCs in series for the dsDNA handles and ssRNA regions. The unfolded Δ*L*_c_^MFE^ of each intermediate conformation was obtained from the fitted *L_c_* value of the ssRNA segment in that conformer, after subtracting the *L*_c_ of the reference MFE structure (see Supplementary Fig. S4 for a detailed description). To determine whether the single-stranded *HTT* exon 1 nucleotides flanking the CAG region contributed to the observed low force transitions, the overhangs of the dsDNA handles were extended to hybridize with these nucleotides. Force ramp experiments using these extended dsDNA handles showed similar results compared to the dsDNA handles with shorter overhangs (Supplementary Fig. S5).

The force associated with each intermediate transition in each FEC was determined using a custom python script that determines the point at which the observed FEC deviates consistently from its fitted eWLC for the folded mRNA. Briefly, all identifiable intermediates in the folded region of each FEC were fit as described above, and the root-mean-squared error (RMSE) of the fitted eWLC from the experimental FEC was calculated using a sliding window. The unfolding force was defined as the first point at which the RMSE permanently crossed a threshold set to the mean + 3σ of the RMSE before the transition (Supplementary Figure S2).

The cooperativity of the main unfolding transition for a subset of FECs of f*HTT*_40_ was defined by the deviation in d*F*/d*x* following the start of the transition. <d*F*/d*x>* and the associated standard deviation (σ) were defined over a slice of the FEC used for fitting the eWLC up to the start of the transition. Then, d*F*/d*x* was evaluated over a window starting from the fitted slice and extending 5 nm past the transition. Transitions were classified as cooperative when d*F*/d*x* in this region dropped below a threshold equal to <d*F*/d*x*> – 3.5σ (Supplementary Figure S3A). Transitions were classified as non-cooperative when d*F*/d*x* failed to drop below this threshold within this region (Supplementary Figure S3B).

### Passive mode assays and HMM fitting

After collecting FECs for a single tethered molecule, a pretension was set at low (∼ 5 pN) and high (∼12 pN) forces corresponding to the transitions observed in the FEC. Fluctuations in the force and extension at constant trap separation (‘passive mode’, PM) were recorded for ∼ 5 min at each pretension before manually relaxing and stretching the tethered RNA. Single trajectories of force versus time, *F*(*t*), were converted to unfolded contour length relative to the MFE structure, Δ*L_c_*^MFE^(*t*), using the eWLC equation described above and the parameter_trace function from the Lumicks Pylake package.

For the low force region of the FECs corresponding to slipped transitions, seven PM traces were collected from five different molecules with pretensions between 4.5 and 9 pN. Following conversion of the data into Δ*L_c_*^MFE^(*t*), traces were concatenated and fit using a step-finding HMM algorithm ^47^. Processed data were first fit to a maximum likelihood step finding algorithm. The data were clustered using a Gaussian Mixture Model (GMM) and the number of states was increased by one until the Bayesian information criterion (BIC) of the model reached a minimum ^45^. The results of step finding and GMM clustering were then used for parameter initialization and fitting of the HMM model. Data were initially fit to a blind HMM to determine the number of states and their positions, yielding a model with 10 states whose distributions of dwell times can be described well by a single exponential function (Supplementary Fig. S10). A second model in which the separation between states was set to one CAG triplet (1.77 nm) agreed well with the blind HMM (Fig. 3C-D, Table S3). PM traces at higher force, near the unfolding transition, were collected from four molecules at a pretension of ∼ 11.8 pN and fitted as described above (Supplementary Fig. S11). The dwell time distributions for each state were fitted to a single exponential function by maximum likelihood estimation to extract the lifetime using the “dwelltimes” module from the Lumicks Pylake package. State means and lifetimes of all models are reported in Supplementary Tables 3 and 4.

### Data availability

All data are available from the lead author upon reasonable request.

## Acknowledgements

This work was supported by grants from the National Institutes of Health (R21NS128701 and T32GM080189 for support of BMO.), and the Pew Innovation Fund.

## Notes

### Competing Interest Statement

The authors have declared no competing interest.

