## Supplementary Information for "Stick-slip unfolding favors self-association of expanded *HTT* mRNA"

Supplementary Figures 1-11

Supplementary Tables 1-4

Supplementary References

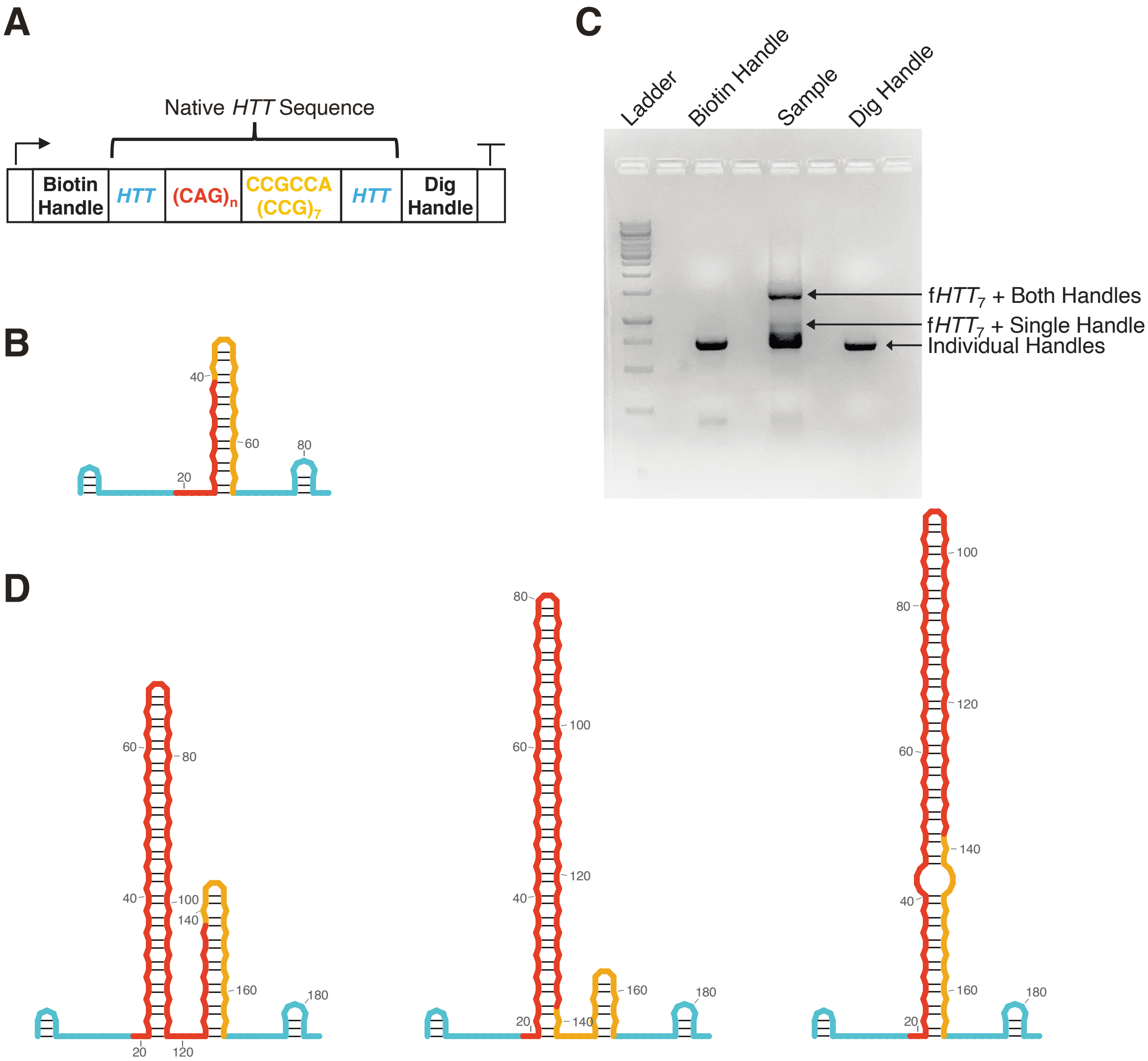


**Supplementary Figure 1. Preparation of tethered HTT RNA. A.** Diagram of transcription template used to generate fragments of HTT exon 1 (f*HTT*$_n$) mRNAs (top). The RNAs used in this study contained n = 7, 12, 20 or 40 CAG repeats (red) plus flanking CCG triplets (gold), additional ~20 HTT residues flanking the repeat region (cyan), and extensions complementary to each DNA handle (black). **B.** MFE structure of the f*HTT*$_7$ mRNA used in this study. The small hairpins predicted in the flanking region (cyan) do not contribute to the observed FECs. **C.** f*HTT* mRNA was refolded and annealed to dsDNA handles, and the product was resolved on a 1.5 % agarose gel with1 kbp dsDNA ladder. **D.** The three most stable secondary structures for f*HTT*$_{40}$ mRNA at 37 ˚C predicted by Mfold [1], colored as in A. The CCG triplets can pair with nearby CAG triplets (left), with themselves (center), or with upstream CAG triplets (right).

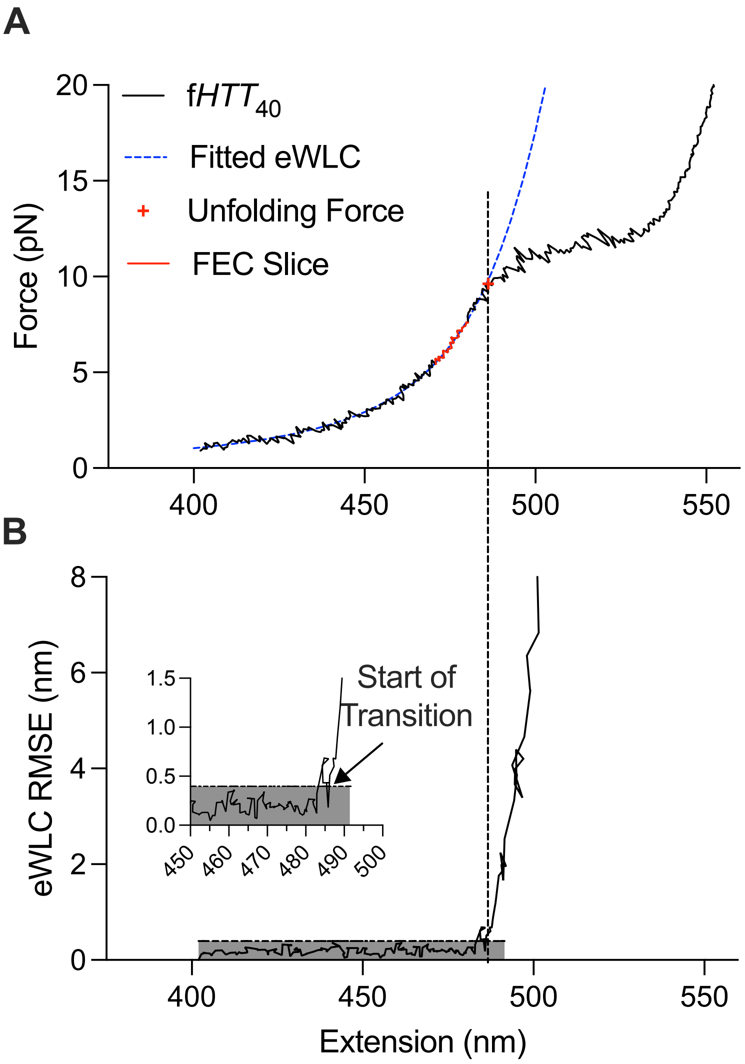


**Supplementary Figure S2. Detection and assignment of unfolding forces.** The unfolding force, $F_U$, was defined by departure of the observed FEC from the predicted elastic behavior of the folded RNA plus DNA handles. See Methods for details. A. FEC from f*HTT*$_{40}$ highlighting the sliced region of the FEC (red line) used to obtain the fitted eWLC (blue dashed line) representing the RNA before unfolding. B. Root mean squared error (RMSE) of the eWLC fitted to the FEC. A noise threshold was defined as equal to the mean RMSE + 3σ, calculated over the FEC slice (gray box). The unfolding force $F_U$ is defined as the first point that crosses this threshold, only if all subsequent points remain outside of the threshold (the point of no return). Inset: Enlarged view of the RMSE permanently crossing the threshold at the start of the unfolding transition.

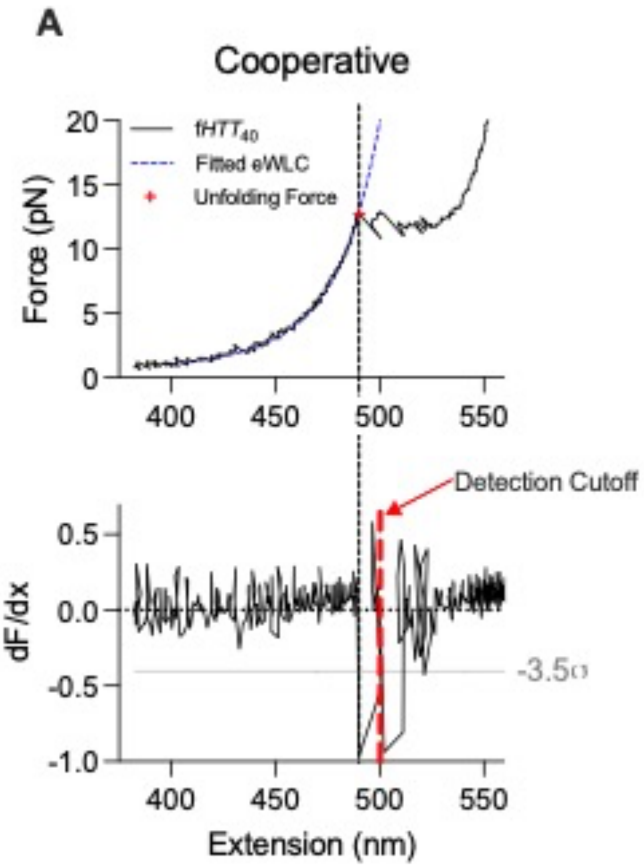


**Supplementary Figure S3. Classification of cooperative vs noncooperative transitions.** A. Top, example of a stretch FEC displaying a cooperative unfolding transition. The unfolding force, $F_U$, was defined as shown in Supplementary Figure 2. Bottom, a transition is cooperative if the derivative of the FEC, d*F*/d*x*, crosses below a threshold equal to <d*F*/d*x*> – 3.5σ, in which σ is the standard deviation of d*F*/d*x* within the analyzed slice up to a detection boundary 5 nm beyond the unfolding event (red dashed lines). B. Top, example of a stretch FEC displaying a noncooperative unfolding transition, as in A. Bottom, noncooperative transitions are defined as FECs for which dF/dx does **not** cross below the threshold within 10 nm of the start of unfolding.

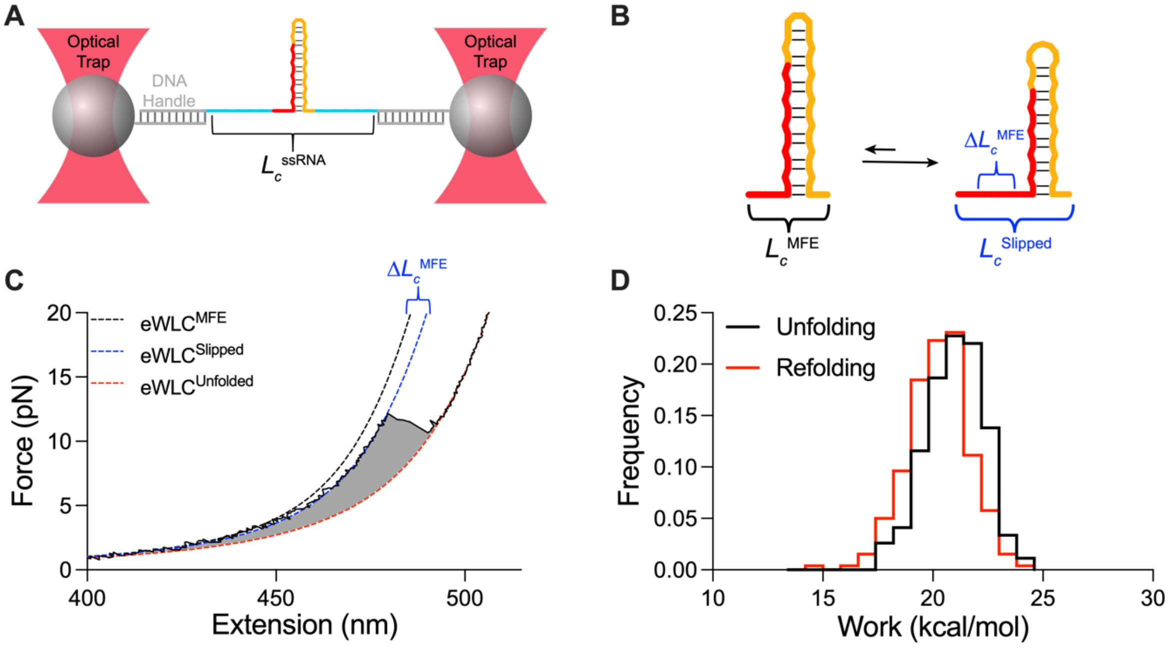


**Supplementary Figure S4. Quantifying folded f*HTT* intermediates.** A. Cartoon of the eWLC model for the folded f*HTT*$_7$ MFE reference structure annealed to dsDNA handles, in which $L_c^{ssRNA}$ represents the end-to-end distance of the first and last ssRNA nucleotides plus one hairpin (2.2 nm; [2]). The RNA 5′ and 3′ ends are annealed to 29-30 nt single-strand extensions of the DNA handles. B. Cartoon illustrating the expected contour lengths for slipped conformations. Each slipped triplet adds three nucleotides to the single-stranded region, increasing its contour length by $L_C$(ss) = 0.59 nm/nt (1.77 nm per triplet). The change in contour length relative to that of the MFE structure ($\Delta L_c^{MFE}$) associated with a slipped transition was determined by fitting the intermediate regions in each FEC and subtracting the fitted $L_c^{ssRNA}$ from the expected $L_c^{ssRNA}$ of the MFE structure (see Methods). C. Example of eWLC fits to the fully folded (black), slipped (blue), and unfolded (red) regions of a force extension curve for f*HTT*$_7$ mRNA. The work associated with the main unfolding and refolding transition (high force peak in Fig. 1C) was calculated by numerical integration of the folded eWLC fitted model up to the end of the unfolding event, and subtraction of the work associated with stretching the dsDNA handles and unfolded RNA over the same window (shaded region). D. The overlap of unfolding and refolding force distributions indicated that the system is near equilibrium. The average work associated with the main transition was ± 20 kcal/mol.

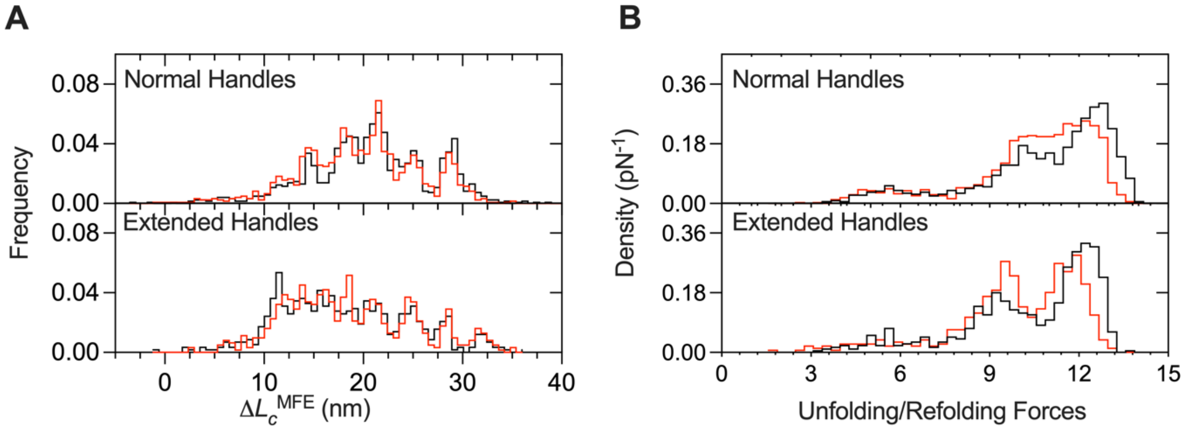


**Supplementary Figure S5. Adjacent *HTT* RNA does not cause intermediate folding.** The dsDNA handles were extended to mask single-stranded $HTT_{20}$ exon1 nucleotides flanking the composite hairpin structure. A-B. Distributions of slipped intermediate contour lengths and unfolding and refolding forces for normal and extended handles. The similar distributions suggest these nucleotides do not form structures that contribute to the observed results.

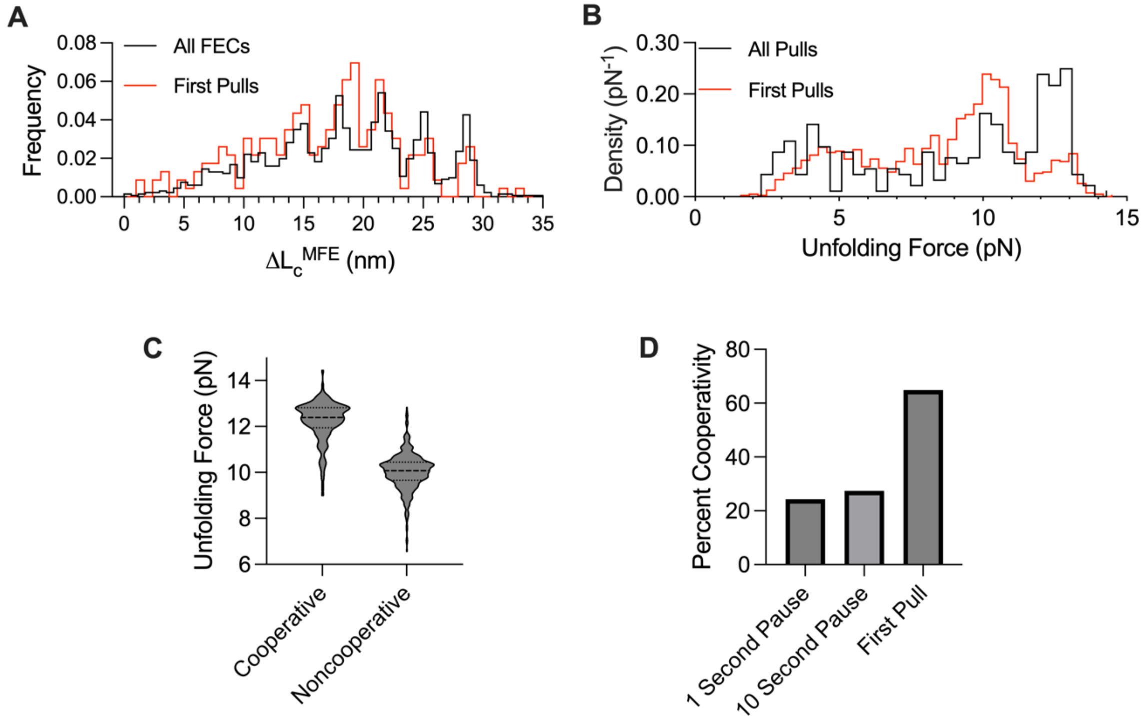


**Supplementary Figure S6**. **Frustrated refolding of expHTT mRNA.** Comparison of first pulls (red) with all FECs (black) for f*HTT*40. A. Distribution of $\Delta L_c^{MFE}$ (intermediate contour lengths) as in Fig. 2B, last row. B. Distributions of unfolding forces. C**.** Distribution of $F_U$ during force-ramp cycles with 1 s equilibration time for f*HTT*40 mRNAs showing that transitions classified as cooperative unfold at higher force (mean = 12.5 pN; *N* = 500 FECs from 99 molecules) than non-cooperative unfolding transitions (mean = 10 pN), on average. Heavy dashed line, median; light dashed lines, 1st and 3rd quartiles. D**.** Fraction of cooperative transitions in f*HTT*40 mRNAs with 1 or 10 s equilibration between pulls, for the first pull. Before the first stretch, molecules were thermally renatured and equilibrated. First pulls, *N* = 95 molecules; 10 s pause, *N* = 294 FECs from 34 molecules.

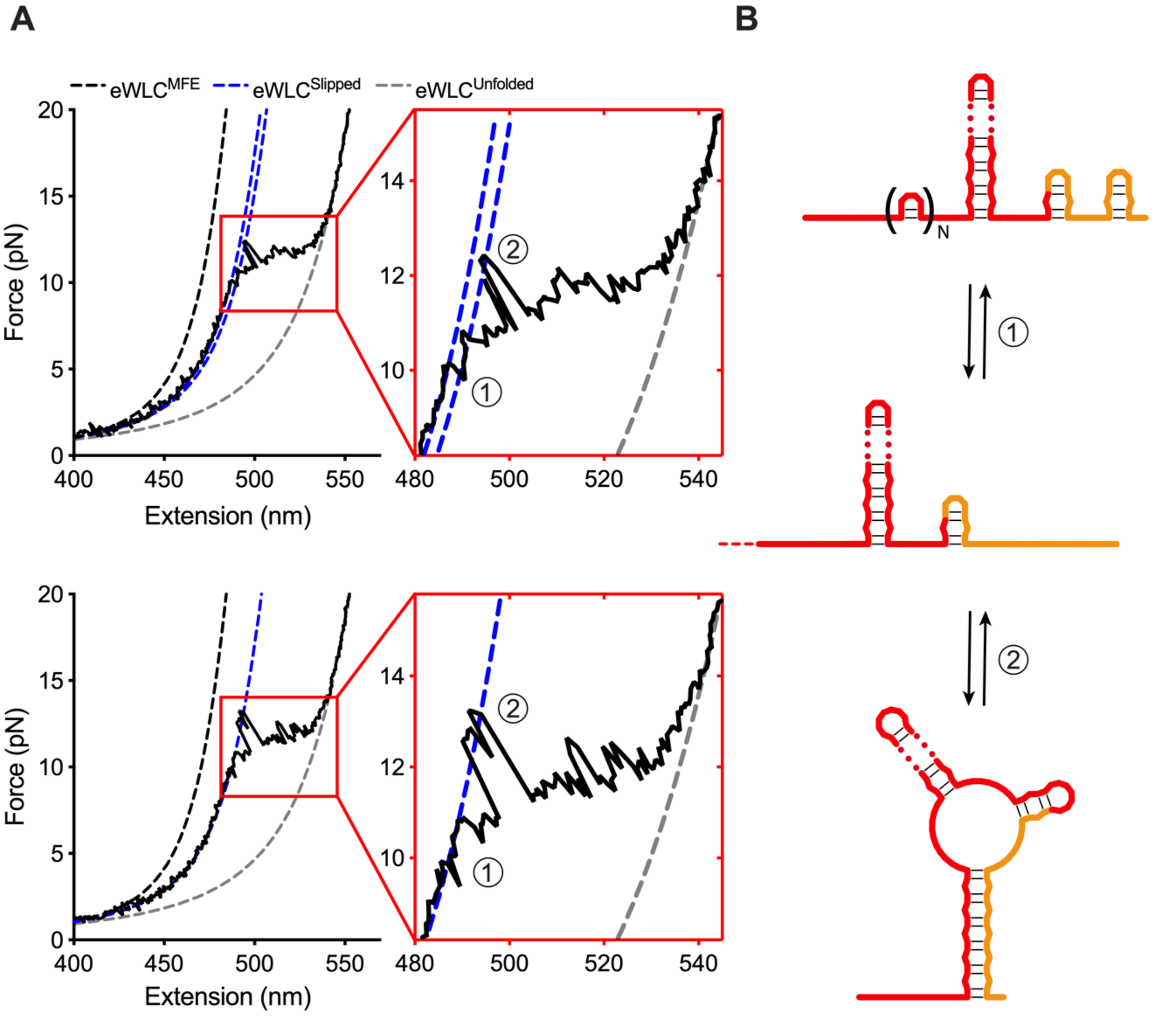


**Supplementary Figure S7. Applied force facilitates base pair exchange.** A. FECs and eWLC fits from two different f*HTT*$_{40}$ molecules after the first pull of a force-ramp cycle. Insets highlight conformational dynamics within the main transition, with many smaller transitions occurring as the RNA gradually extends (1). Occasionally, the tethered *fHTT* mRNA transitions to a much higher force (2) just after the start of a non-cooperative transition (1), reflecting temporary refolding to a structure with a shorter contour length. B. Cartoon depicting how applied force induces base pair shuffling that exposes both CAG and CCG ssRNA triplets. The exposed bases can participate in transient long range base pairs with the flanking CCG repeats.

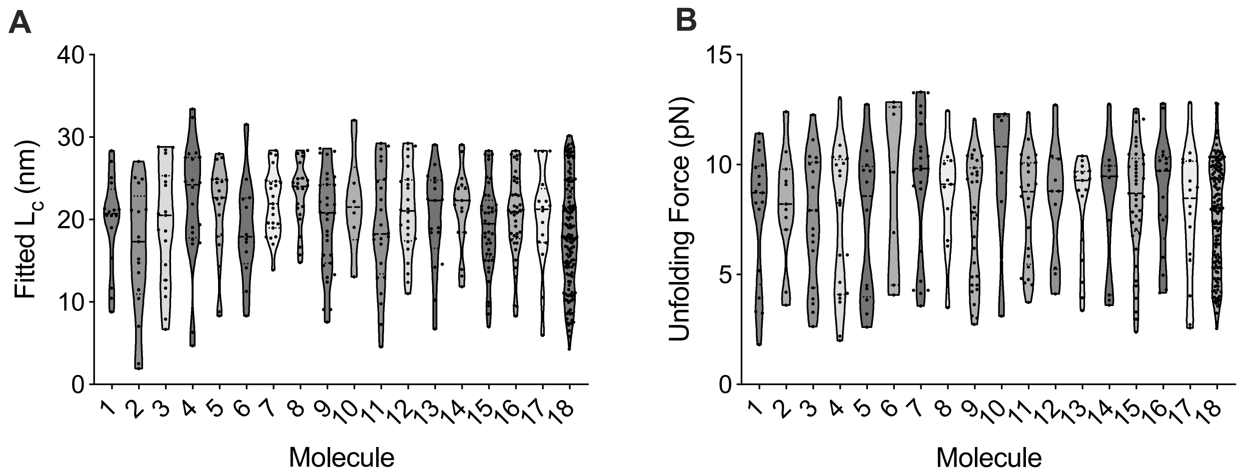


**Supplementary Figure S8**. **Variable unfolding of single f*HTT*$_{40}$ molecules.** A. Violin plots of fitted intermediate contour lengths for all FECs from 18 of 95 molecules used in this study. Solid line, median; dashed lines, upper and lower quartile. Each molecule visited different intermediates during successive force-ramp cycles. B. Same as (A) but for unfolding forces.

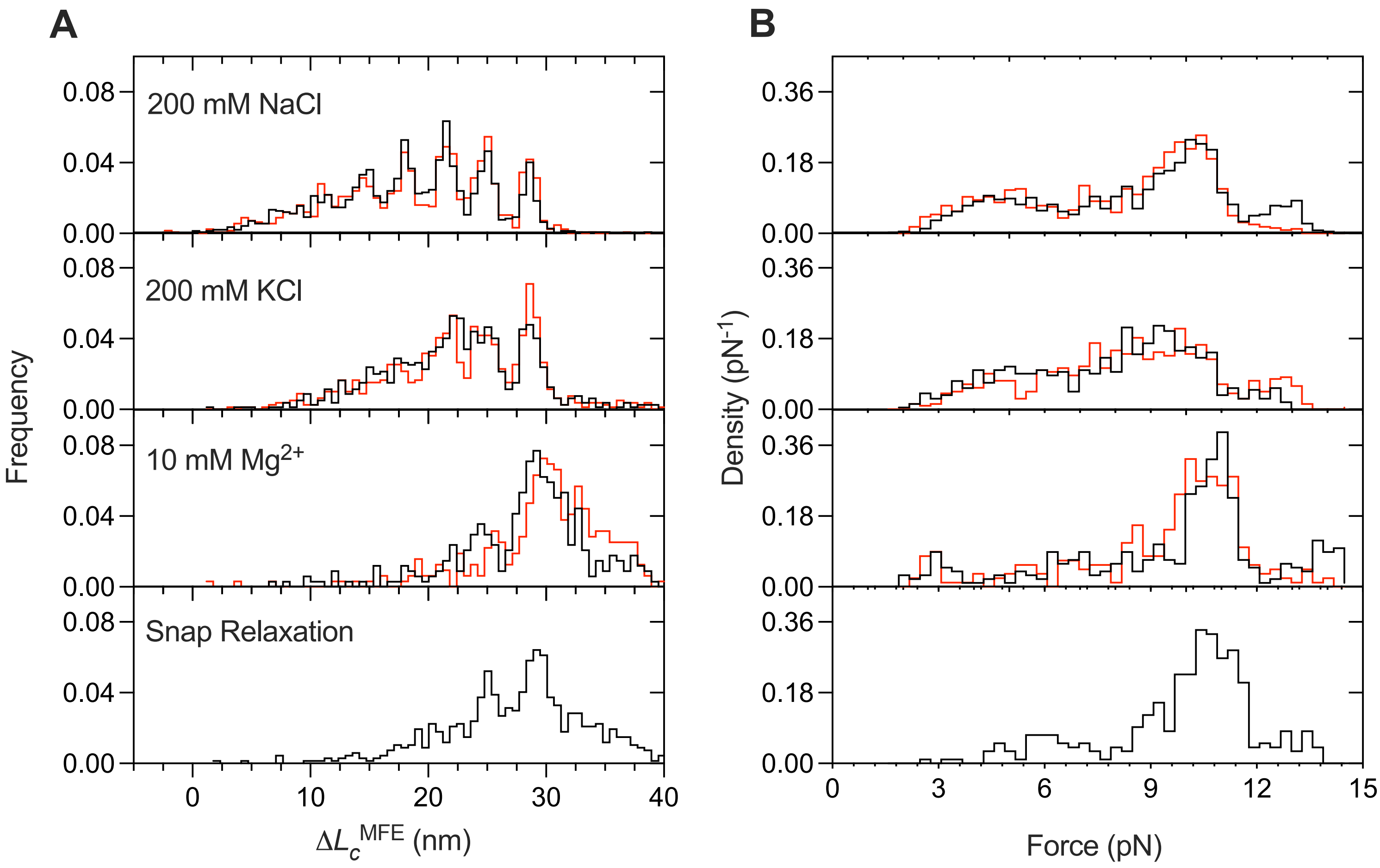


**Supplementary Figure S9**. **Folding heterogeneity in different experimental conditions.** Force ramp experiments were performed with f*HTT*$_{40}$ in the standard assay buffer but with the counterions shown. For snap relaxation, tethers were stretched at 90 nm/s followed by a rapid relaxation at 500 nm/s. Only stretch FECs were analyzed for this condition ($N$ = 203 FECs from 11 molecules). For all histograms, black and red lines indicate stretches and relaxations, respectively. A. Intermediate contour lengths. B. Unfolding and refolding forces.

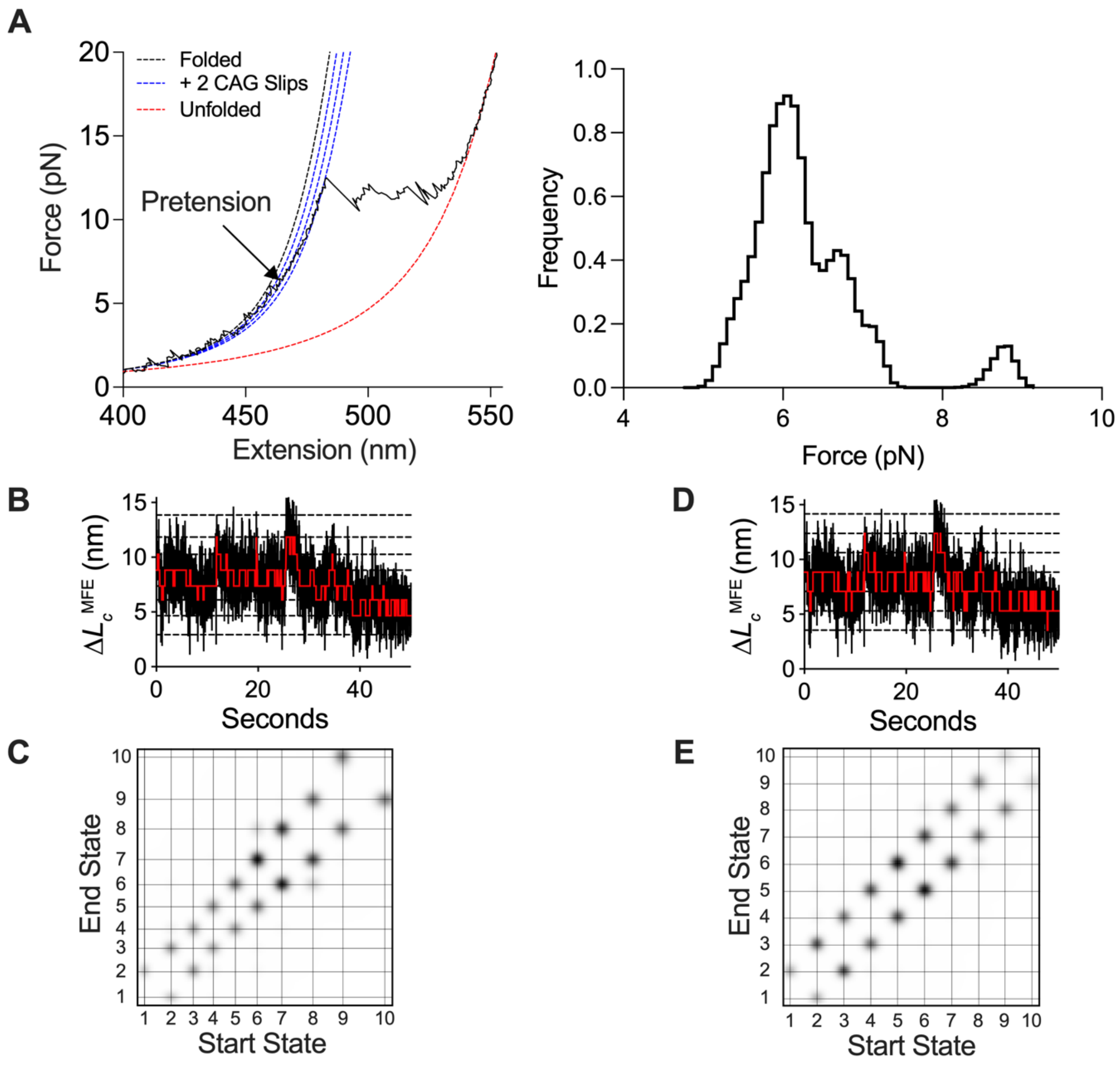


**Supplementary Figure S10. Passive mode assay to measure the slipping kinetics at low force.** A. Representative *fHTT*$_{40}$ FEC indicating the applied pretension, which was below the main unfolding transition (right). Left, force distribution during passive mode (PM) observation at 7 pretensions from 4.5 to 9 pN. $F(t)$ vs. $t$ recordings at each pre-tension were concatenated before further analysis. B-C. Results of blind HMM fit to low force PM trace (B) and transition density plot (C). The best fit (BIC) was obtained with 10 states, from less extended to more extended as shown. The results of the blind model were used to assign fixed states. D-E. Results of fixed HMM fit to PM trace (D) and transition density plot. (E). HMM was fitted to the same data as in B,C. See Table S3 for detailed results. See methods for HMM fitting details.

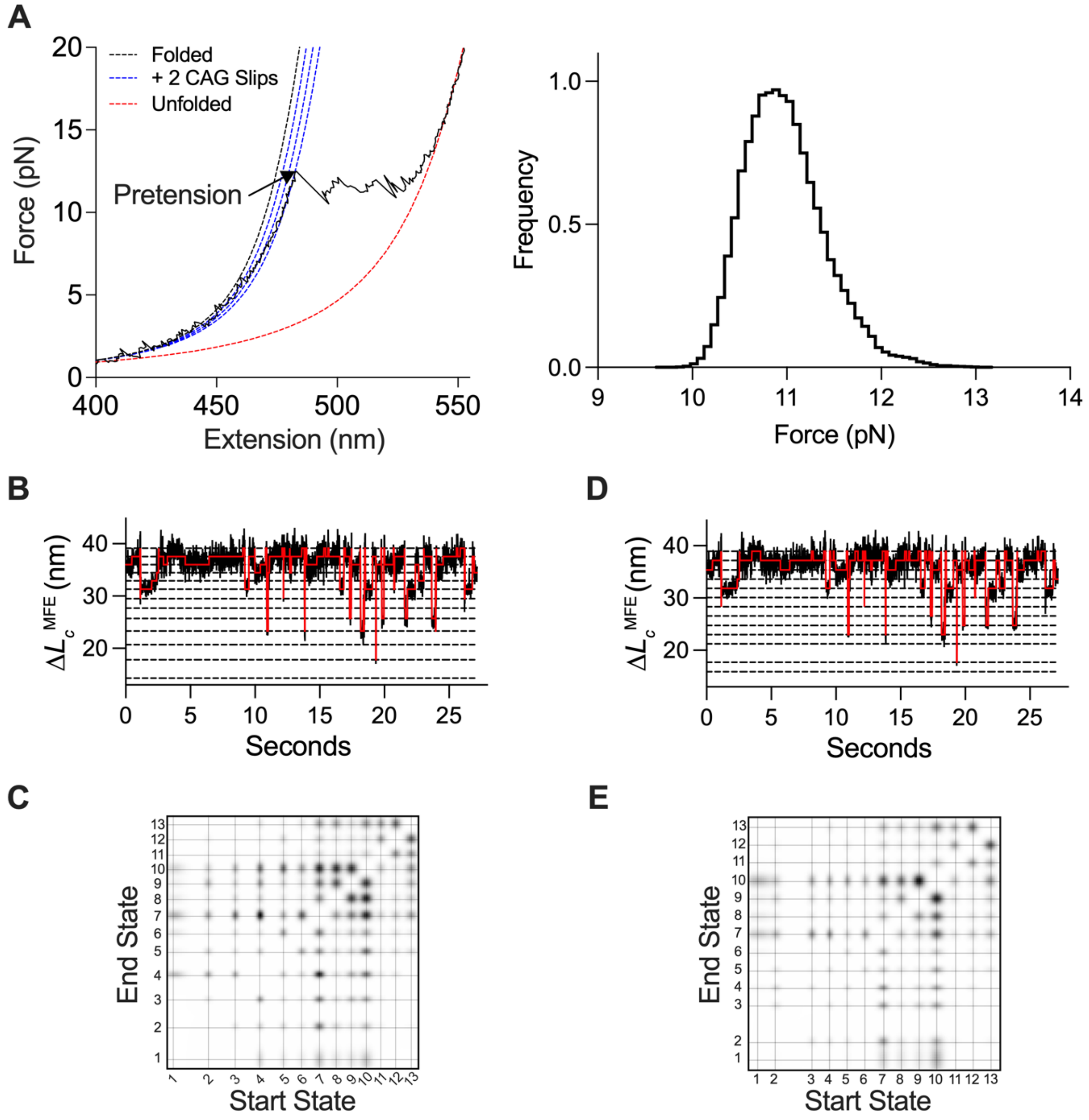


**Supplementary Figure S11**. **Passive mode analysis of folding kinetics at high force.** Data acquisition and analysis as in Fig. S10. The pretension was set near the start of the main unfolding transition. HMM was fitted to four PM traces obtained from four molecules. See Table S4 for detailed results.

### Supplementary Tables

**Table S1. Primers to generate dsDNA handles[a] and f*HTT* transcription templates[b]**

| Name | Sequence |
|---|---|
| **Biotin Handle Fwd Primer** | 5′ /**5B**/ CCGCTTACAGACAAGCTGTGACCGTCTCC 3' |
| **Biotin Handle Rev Primer** | 5′CTCTCTACCCATCCATCTCTCACTCACCC/**Ab**/GGAGAAAATGAAGACCGCGACGGGTGTC 3' |
| **Biotin Handle Rev Primer (Extended)** | 5′AGGACTTGAGGGACTCCTCTCTACCCATCCATCTCTCACTCACCC/**Ab**/GGAGAAAATGAAGACCGCGACGGGTGTC 3' |
| **Dig Handle Fwd Primer** | 5′CACTTTGGACAGGACACACAGGACACAGG/**dU**/GGTTTTCACCGTCATCACCGAAACG 3' |
| **Dig Handle Fwd Primer (Extended)** | 5′CCTCAGCTTCCTCAGCCTCACTTTGGACAGGACACACAGGACACAGG/**dU**/GGTTTTCACCGTCATCACCGAAACG3' |
| **Dig Handle Rev Primer** | 5′/**DigN**/GGAGAAAATGAAGACCGCGACGGGTGTCTGGTCAGAGACATC 3' |
| **f*HTT* Transcription Template[c]** | 5′GGATCCTAATACGACTCACTATAGGGGGTGAGTGAGAGATGGATGGGTAGAGAGGAGTCCCTCAAGTCCTTCCAGCAGCAGCAGCAGCAGCAGCCGCCACCGCCGCCGCCGCCGCCGCCGCCTCCTCAGCTTCCTCAGCCTCACTTTGGACAGGACACACAGGACACAGGCTAGCATAACCCCTTGGGGCCTCTAAACGGGTCTTGAGGGGTTTTTTGGCATGC 3' |
| **Transcription Template Fwd Primer** | 5′TAATACGACTCACTATAGGGGGTGAGTGAGAGATGGATGGGTAG3' |
| **Transcription Template Rev Primer** | 5′CAAAAAACCCCTCAAGACCCGTTTAGAGGCCCCAAGGGGTTATGCTAGCCTGTGTCCTGTGTGTCCTGTCC 3' |

[a]Primer sequences used to generate dsDNA handles. Bold insertions indicate modifications: **5B** – 5′ Biotin; **Ab** – Abasic site; **dU** – 2′ deoxyuridine; **5DigN** – 5′ digoxygenin.

[b]DNA template used to transcribe f*HTT* mRNAs. T7 promoter and terminator sequences are highlighted in gray. Nucleotides complementary to the overhangs of the dsDNA handles are shown in green. *HTT* exon 1 nucleotides flanking the repeat are highlighted in cyan. The CAG and CCG repeat tracts are highlighted in red and yellow, respectively. Primers used to amplify transcription template shown below.

[c]Number of CAG triplets was $n$ = 7, 12, 20, 40

**Table S2. Populations of f*HTT*$_7$ slipped intermediates relative to $\Delta L_c^{MFE}$.**[a]
Related to Figure 1C.

| State | Amplitude | μ (nm) | SD (nm) |
|---|---|---|---|
| **1** | 0.0455 ± 0.00201 | -1.76 ± 0.036 | 0.58 ± 0.014 |
| **2** | 0.0380 ± 0.00246 | 0.187 ± 0.061 | 0.58 ± 0.014 |
| **3** | 0.0152 ± 0.00253 | 1.55 ± 0.145 | 0.58 ± 0.014 |
| **4** | 0.111 ± 0.00224 | 3.87 ± 0.017 | 0.58 ± 0.014 |
| **5** | 0.0235 ± 0.00266 | 5.65 ± 0.129 | 0.58 ± 0.014 |
| **6** | 0.00487 ± 0.00296 | 6.92 ± 0.511 | 0.58 ± 0.014 |

[a]Parameters obtained from a non-linear least squares regression fit of the distribution of $\Delta L_c^{MFE}$ obtained for f*HTT*$_7$ mRNA to a sum of 6 Gaussians, representing different slipped intermediates. Each state shows the amplitude (relative population), mean (μ) value of $\Delta L_c^{MFE}$ and standard deviation (SD) ± SE obtained from the fit. All states were assumed to have a single SD value to ensure equal widths. Results were obtained from 286 FECs from 48 different molecules. Fit parameters and errors were obtained from the GraphPad Prism software.

**Table S3. HMM parameters for f*HTT*$_{40}$ mRNA transitions at low tension.**[a] Related to Figure 3 and Figure S9.

| State | Free HMM[b] <$\Delta L_c^{MFE}$> ± SE (nm) | Free HMM[b] Lifetime (95% CI, s) | Fixed HMM[c] <$\Delta L_c^{MFE}$> ± SE (nm) | Fixed HMM[c] Lifetime (95% CI, s) |
|---|---|---|---|---|
| 1 | 2.931 ± 0.013 | 0.897 (0.667, 1.247) | 3.54 ± 0.012 | 1.134 (0.873, 1.509) |
| 2 | 4.659 ± 0.016 | 0.579 (0.485, 0.698) | 5.31 ± 0.013 | 0.661 (0.573, 0.768) |
| 3 | 6.112 ± 0.016 | 1.002 (0.85, 1.194) | 7.08 ± 0.013 | 0.852 (0.748, 0.976) |
| 4 | 7.369 ± 0.017 | 0.954 (0.814, 1.128) | 8.85 ± 0.014 | 0.993 (0.872, 1.137) |
| 5 | 8.818 ± 0.017 | 1.12 (0.967, 1.306) | 10.62 ± 0.014 | 0.712 (0.636, 0.801) |
| 6 | 10.263 ± 0.017 | 0.597 (0.534, 0.669) | 12.39 ± 0.014 | 0.358 (0.32, 0.401) |
| 7 | 11.857 ± 0.018 | 0.392 (0.352, 0.439) | 14.16 ± 0.014 | 0.443 (0.388, 0.51) |
| 8 | 13.847 ± 0.018 | 0.378 (0.336, 0.427) | 15.93 ± 0.014 | 0.43 (0.371, 0.503) |
| 9 | 15.745 ± 0.019 | 0.47 (0.41, 0.543) | 17.7 ± 0.015 | 0.247 (0.204, 0.304) |
| 10 | 18.553 ± 0.022 | 0.306 (0.251, 0.379) | 19.47 ± 0.017 | 0.392 (0.27, 0.602) |

[a]Mean $\Delta L_c^{MFE}$ and lifetime for each conformational state, obtained from HMM fits to F(t). Seven passive mode traces were collected for five molecules of f*HTT*$_{40}$ mRNA at low pretensions (5-9 pN). The HMM was fit using the sfHMM Python package from [S2]. The standard errors of each state mean were determined by the covariance matrix of the HMM fit. Lifetimes were obtained by single exponential fits to the distribution of dwell times for each state. Lifetime fitting and confidence intervals were determined using the Lumicks Pylake Python package. See Methods for details.

[b]The number of states and their means were determined by BIC minimization.

[c]Mean $\Delta L_c^{MFE}$ values were fixed and separated by the length of one single-stranded CAG triplet (~1.77 nm).

**Table S4. HMM parameters for f*HTT*$_{40}$ mRNA transitions at high tension[a].** Related to Figure 3 and Figure S10

| State | Free HMM $<\Delta L_c^{MFE}> \pm$ SE (nm) | Free HMM Lifetime (95% CI, s) | Fixed HMM $<\Delta L_c^{MFE}> \pm$ SE (nm) | Fixed HMM Lifetime (95% CI, s) |
|---|---|---|---|---|
| 1 | 14.291 ± 0.098 | 0.061 (0.049, 0.077) | 15.93 ± 0.127 | 0.058 (0.047, 0.073) |
| 2 | 17.851 ± 0.041 | 0.093 (0.078, 0.112) | 17.7 ± 0.042 | 0.109 (0.091, 0.131) |
| 3 | 20.7 ± 0.035 | 0.151 (0.129, 0.18) | 21.24 ± 0.036 | 0.157 (0.133, 0.187) |
| 4 | 23.311 ± 0.036 | 0.093 (0.083, 0.106) | 23.01 ± 0.03 | 0.111 (0.095, 0.13) |
| 5 | 25.738 ± 0.038 | 0.156 (0.135, 0.182) | 24.78 ± 0.032 | 0.127 (0.107, 0.153) |
| 6 | 27.696 ± 0.045 | 0.386 (0.331, 0.455) | 26.55 ± 0.036 | 0.332 (0.275, 0.405) |
| 7 | 29.53 ± 0.05 | 0.151 (0.138, 0.166) | 28.32 ± 0.048 | 0.169 (0.154, 0.187) |
| 8 | 31.311 ± 0.054 | 0.429 (0.378, 0.489) | 30.09 ± 0.047 | 0.311 (0.273, 0.357) |
| 9 | 32.905 ± 0.055 | 0.505 (0.449, 0.571) | 31.86 ± 0.053 | 0.446 (0.398, 0.503) |
| 10 | 34.448 ± 0.058 | 0.241 (0.219, 0.265) | 33.63 ± 0.058 | 0.306 (0.282, 0.334) |
| 11 | 35.988 ± 0.045 | 0.52 (0.434, 0.63) | 35.4 ± 0.044 | 0.437 (0.371, 0.519) |
| 12 | 37.555 ± 0.05 | 0.57 (0.483, 0.68) | 37.17 ± 0.047 | 0.509 (0.44, 0.593) |
| 13 | 39.172 ± 0.054 | 0.061 (0.053, 0.07) | 38.94 ± 0.051 | 0.083 (0.073, 0.094) |

[a]Mean $\Delta L_c^{MFE}$ and lifetimes obtained from HMM fits to four passive mode traces from four molecules of f*HTT*$_{40}$ mRNA at high pretension (~12 pN). State means, lifetimes and errors were determined as in Table S3.
